# Single-molecule reaction chambers reveal intramolecular protein kinase activation

**DOI:** 10.64898/2026.10.08.757607

**Authors:** Eva Krötenheerdt, Alina Ehrl, Christian Stelmach, Francesco Reina, Lucas Piëch, Florian Praetorius, Jonas Ries, Bojan Žagrović, Thomas A. Leonard

## Abstract

Many protein kinases, ubiquitous signal transducers across all kingdoms of life, are activated by phosphorylation of one or more conserved residues in their activation loops. This reaction is widely considered to be intermolecular (*trans*) in nature, dependent on the transient dimerization of two kinase domains. However, since the substrate, product and reaction chemistry are identical, bulk solution kinase assays cannot distinguish between intra- and inter-molecular reactions. Discriminating between these two mechanisms is crucial because they impose different requirements on the cell for acute, high-fidelity signal transduction. Using single-molecule reaction chambers fabricated from DNA origami, we show that the paradigmatic insulin receptor kinase (IRK) undergoes intramolecular (*cis*) autophosphorylation with kinetics indistinguishable from those in free solution. Furthermore, we demonstrate that the bacterial serine/threonine kinase protein kinase B (PknB) undergoes intramolecular autophosphorylation, allosterically stimulated by intermolecular (*trans*) interactions. Our findings establish that intramolecular activation loop autophosphorylation is not only possible, but likely an ancient and conserved feature of protein kinase signaling. More broadly, our work provides new tools for single-molecule enzymology.

## Introduction

Protein phosphorylation is a ubiquitous and central regulator of cellular signal transduction across all kingdoms of life^1^. Since the discovery of protein kinases in 1955^2^, considerable effort has been devoted to understanding their molecular regulation. Kinases constitute a large protein family likely derived from a common ancestral kinase^1,3^. A recurring feature of the activation of many kinases is the phosphorylation of a conserved residue in the activation loop^4^. However, despite decades of study, the origin and mechanism of this modification remains controversial^5,6^.

Conceptually, activation loop autophosphorylation can proceed through two distinct mechanisms: (1) in an intramolecular *cis*-autophosphorylation reaction, a kinase catalyzes the phosphorylation of its own activation loop; (2) in an intermolecular *trans* reaction, protomers of a dimeric kinase reciprocally phosphorylate the activation loop of their opposing protomer. Whilst chemically identical, *cis-*and *trans*-autophosphorylation are fundamentally different modes of regulation, not a subtle semantic nuance. A *trans* reaction requires kinase molecules to encounter one another within the crowded cellular environment in a timely manner and with the correct geometry for catalysis^7^. An unphosphorylated kinase must be capable of autophosphorylation, but inactive against non-self substrates, a paradox commonly known as the ‘enigma of autophosphorylation^6^.’ Intermolecular activation loop phosphorylation necessitates that dimerization is both face-to-face and transient, such that, following phospho-transfer, the two kinase domains dissociate to permit the binding of downstream substrates. In contrast, a *cis* mechanism requires a distinct autoinhibited state^8^ that prevents constitutive autoactivation, and a mechanism to convert it to an active one upon pathway stimulation.

Our mechanistic understanding of kinase activation by activation loop autophosphorylation has been heavily influenced by the structure of IRK^9^, one of the earliest protein kinase domain structures determined. The structure revealed the activation loop of IRK seemingly poised for intramolecular phospho-transfer, but in a conformation sterically incompatible with ATP binding. It was thus inferred that the activation loop conformation was autoinhibitory and that intramolecular autophosphorylation was not possible. Instead, activation loop phosphorylation was proposed to be catalyzed intermolecularly by a second kinase protomer within the insulin receptor^9^. The model has since been reinforced by the dimeric architecture of the active state of many receptor tyrosine kinases as well as a wealth of kinase domain crystal structures in which activation loop exchange has been visualized between neighboring protomers in the crystal lattice, thereby positioning the activation loop of one protomer within the catalytic cleft of the other^10^. These include structures of the insulin-like growth factor receptor (IGFR) kinase, which exhibit both symmetric and asymmetric activation loop-exchanged dimers in the crystal lattice^11,12^. Although the physiological relevance of some of these crystallographic assemblies remains uncertain^5,10^, collectively these observations established *trans*-autophosphorylation as the canonical mechanism of kinase activation^5,6,10,13–17^. Concentration-dependent kinetics have since been used to infer the intermolecular nature of the autophosphorylation reaction ^18–20^.

A growing body of evidence, however, challenges this paradigm. Several kinases, including the dual-specificity tyrosine-phosphorylation-regulated protein kinases (DYRKs)^21^, glycogen synthase kinase 3β (GSK3β)^22^, extracellular signal-regulated kinase (ERK)^27,28^, mitogen-activated protein kinase (Fus3)^23^, AuroraA (AurA)^20^, Checkpoint kinase 2 (Chk2)^20^, Protein Kinase D (PKD)^24^, the integrated stress response (ISR) kinases^25,26^, and the unfolded protein response (UPR) kinase Inositol-requiring enzyme 1 (IRE1)^25^ have been reported to undergo intramolecular (*cis*) activation loop autophosphorylation. These findings raise an important question: are these kinases genuine exceptions to an otherwise universal mechanism, or is intramolecular autophosphorylation the rule as opposed to the exception?

To address this question, we revisited IRK, the structural archetype upon which the *trans*-autophosphorylation model was originally built. Our molecular dynamics simulations reveal that its activation loop can adopt conformations compatible with intramolecular autophosphorylation, demonstrating that activation loop *cis*-autophosphorylation is sterically feasible. To unambiguously test this prediction experimentally, we built single-molecule reaction chambers out of DNA origami. By encapsulating individual IRK molecules within the reaction chambers, we demonstrate that IRK undergoes intramolecular activation-loop phosphorylation of all three canonical tyrosines. To determine whether *cis*-autophosphorylation is conserved beyond IRK, we studied the bacterial serine/threonine kinase PknB from *Mycobacterium tuberculosis*. We demonstrate that conventional concentration-dependent kinase assays cannot distinguish direct intermolecular phosphorylation from an intramolecular reaction that is allosterically stimulated by intermolecular interactions. Taken together, our findings argue that intramolecular autophosphorylation is likely a widespread mechanism of kinase activation, explain why kinase assays have been systematically interpreted in the framework of *trans*-phosphorylation, and establish that intermolecular regulation of autophosphorylation is mechanistically distinct from intermolecular phospho-transfer.

## Results

### Intramolecular activation loop autophosphorylation is sterically possible

The current model of IRK activation proposes that insulin binding to the ectodomain of the receptor elicits a large conformational change which results in the cytoplasmic kinase domains being brought into close enough proximity such that they can undergo intermolecular *trans*-autophosphorylation on Y1158, Y1162, and Y1163^29^. However, the repositioning of the kinase domains was inferred and has not been directly observed. In principle, the conformational change associated with insulin binding could also elicit, allosterically, intramolecular c*is*-autophosphorylation (Supplementary Figure 1A). The insulin receptor belongs to a sub-family of receptor tyrosine kinases (Supplementary Figure 1B), for which the activation loop adopts a conserved conformation in experimentally determined structures (Supplementary Figure 1C). Although the conformation of the activation loop of IRK has been described to be incompatible with intramolecular autophosphorylation^9^, Y1162 is superimposable with the position of the acceptor tyrosine in the structure of IRK in complex with ATP and a substrate peptide^30^ (Supplementary Figure 1D-E) and it is rather the N-terminal segment of the activation loop which sterically occludes ATP binding^9^. This segment, however, exhibits the highest thermal displacement (B-factors) in the structure of apo-IRK (Supplementary Figure 1F), indicating that its conformational flexibility is high. Together, these observations raise the question of whether the activation loop can sample the conformational space necessary for intramolecular autophosphorylation.

To test this explicitly, we employed classical molecular dynamics (MD) simulations to determine whether each of the activation loop tyrosine residues in IRK (Y1158, Y1162 and Y1163, Supplementary Figure 1G) can be brought sufficiently close to the gamma phosphate of ATP (ATPγP) in the active site to enable intramolecular (*cis*-) autophosphorylation without significantly distorting the rest of the structure. Starting from the inactive structure of the enzyme (1IRK), we used distance and position restraints to bring each individual tyrosine OH group separately into close proximity of ATPγP, followed by unrestrained 1 µs simulations to analyze the stability of phosphorylation competent conformations.

Our simulations show that each of the three activation-loop tyrosines can repeatedly adopt structurally undistorted conformations in which the distance between the tyrosine OH groups and ATPγP is compatible with *cis*-autophosphorylation (Supplementary Figure 1H). Specifically, the observed minimum distance between the oxygen atom of the tyrosine hydroxyl and the phosphorous atom of the gamma phosphate (ATPγP) in unrestrained trajectories was 2.96 Å for Y1158, 4.08 Å for Y1162 and 3.14 Å for Y1163 (Supplementary Figure 1H), which is generally considered compatible with phosphoryl transfer^31^. Y1158 exhibited the most persistent ATPγP -proximal sampling, remaining below 5 Å for 52% of the time. Y1162 and Y1163 also reached the sub-5 Å range multiple times, but in a rather transient manner (Supplementary Figure 1H-I). Importantly, the structure of the kinase domain, excluding the activation loop, remained stable in all simulations, with the average backbone RMSD from the inactive, experimental structure in the 1.5 – 3.0 Å range (Supplementary Figure 1J). Finally, the minimum-distance conformations (Supplementary Figure 1I) as well as the average conformation in unrestrained simulations were, across the board, structurally more similar to the inactive 1IRK structure^9^ than to the active 1IR3 one^30^, as judged by backbone RMSD (Supplementary Figure 1J). However, while MD simulations show that *cis*-autophosphorylation is, in principle, sterically possible, they do not provide evidence that it actually occurs. For this, we need an experimental method to distinguish unambiguously between intra- and inter-molecular reactions.

### Fabrication of single molecule reaction chambers from DNA origami

Both the substrates and products of *cis*- and *trans*-catalyzed reactions are chemically identical. Therefore, bulk solution experiments cannot distinguish between them. To determine whether IRK can phosphorylate its own activation loop intramolecularly, we sought to isolate individual kinase domains in single-molecule reaction chambers made out of DNA origami (Figure 1A).

**Figure 1.**
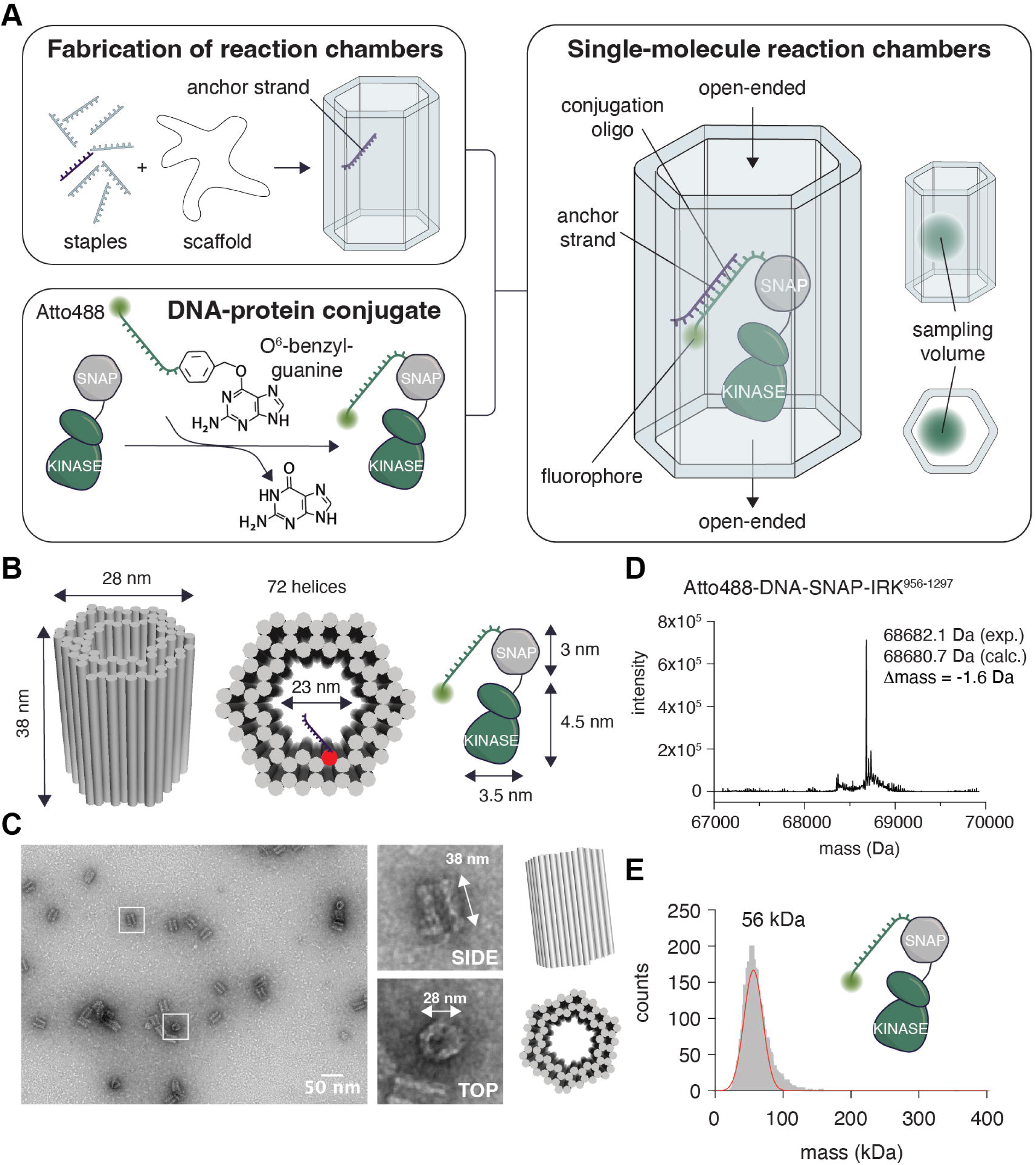
Fabrication of single molecule reaction chambers from DNA origami. A. Schematic depicting the design principles, assembly, and utility of single molecule reaction chambers for biology. The chamber is fabricated using DNA origami, whereas the molecule of interest is fused to the SNAP tag, which can be covalently conjugated with a single-stranded oligonucleotide bearing a 5’ or 3’ benzylguanyl group. For spectroscopic detection and imaging, the oligonucleotide also carries a fluorophore at its other end. B. The chambers comprise 72 helices arranged to form a hexagonal cylinder 38 nm x 28 nm. The inner chamber is large enough to encapsulate a SNAP-kinase domain (SNAP-KD) fusion protein of approximately 3.5 nm x 7.5 nm. C. Negative stain electron microscopy of DNA origami structures. Inset: top and side views of the structures. D. Intact mass spectrometry of the DNA-SNAP-IRK conjugate. E. Mass photometry of the DNA-SNAP-IRK conjugate.

Hexagonal, honeycomb architectures have previously been shown to be suitable for robust self-assembly of DNA origami structures^32^. We therefore designed a hexagonal cylinder with an internal diameter large enough to accommodate our molecule of interest and a centrally located single-stranded anchor strand for target immobilization by complementary base pairing (Figure 1B, Supplementary Figure 2). The cylinder is open-ended, permitting the free exchange of small molecule reaction components. To specifically load the chambers with IRK, we covalently conjugated a SNAP-IRK fusion protein to a single-stranded DNA oligo (conjugation oligo) complementary to the single-stranded overhang of the anchor strand (Figure 1A).

We confirmed the correct assembly and dimensions of the chambers by electron microscopy (Figure 1C) as well as the correct mass and phosphorylation state (unphosphorylated) of the DNA-SNAP-IRK conjugate by mass spectrometry (Figure 1D). Finally, we confirmed that the DNA-SNAP-IRK conjugate is monomeric and monodisperse by mass photometry (Figure 1E). The correct masses of all recombinant protein and DNA-protein conjugates subsequently used in this study were confirmed by mass spectrometry (Supplementary Figure 3). The next step was to immobilize single molecules inside the chambers.

### Reaction chambers can be loaded sequence-specifically

Electrophoretic mobility shift assays (EMSA) with the conjugation oligo show that its incorporation into the origami cylinders is dependent on the presence of its complementary anchor strand (Figure 2A). Similarly, incorporation of the DNA-protein conjugate also requires the presence of the complementary anchor strand. Due to its larger size, however, its incorporation into the cylinders was less efficient than the oligo alone. To determine the molar ratio of cylinders to conjugate that was necessary for stoichiometric loading of the cylinders we performed titration experiments. A ratio of 1:25 cylinders:conjugate was sufficient for stoichiometric loading (Figure 2B). Following stoichiometric loading of the chambers, we removed unbound DNA-protein conjugate by size exclusion chromatography (Figure 2C). For downstream applications requiring the release of the protein of interest from the reaction chamber, we used a toehold oligonucleotide with higher affinity for the conjugation oligonucleotide than the anchor strand to competitively displace the DNA-SNAP-IRK conjugate from the chamber (Supplementary Figure 4).

**Figure 2.**
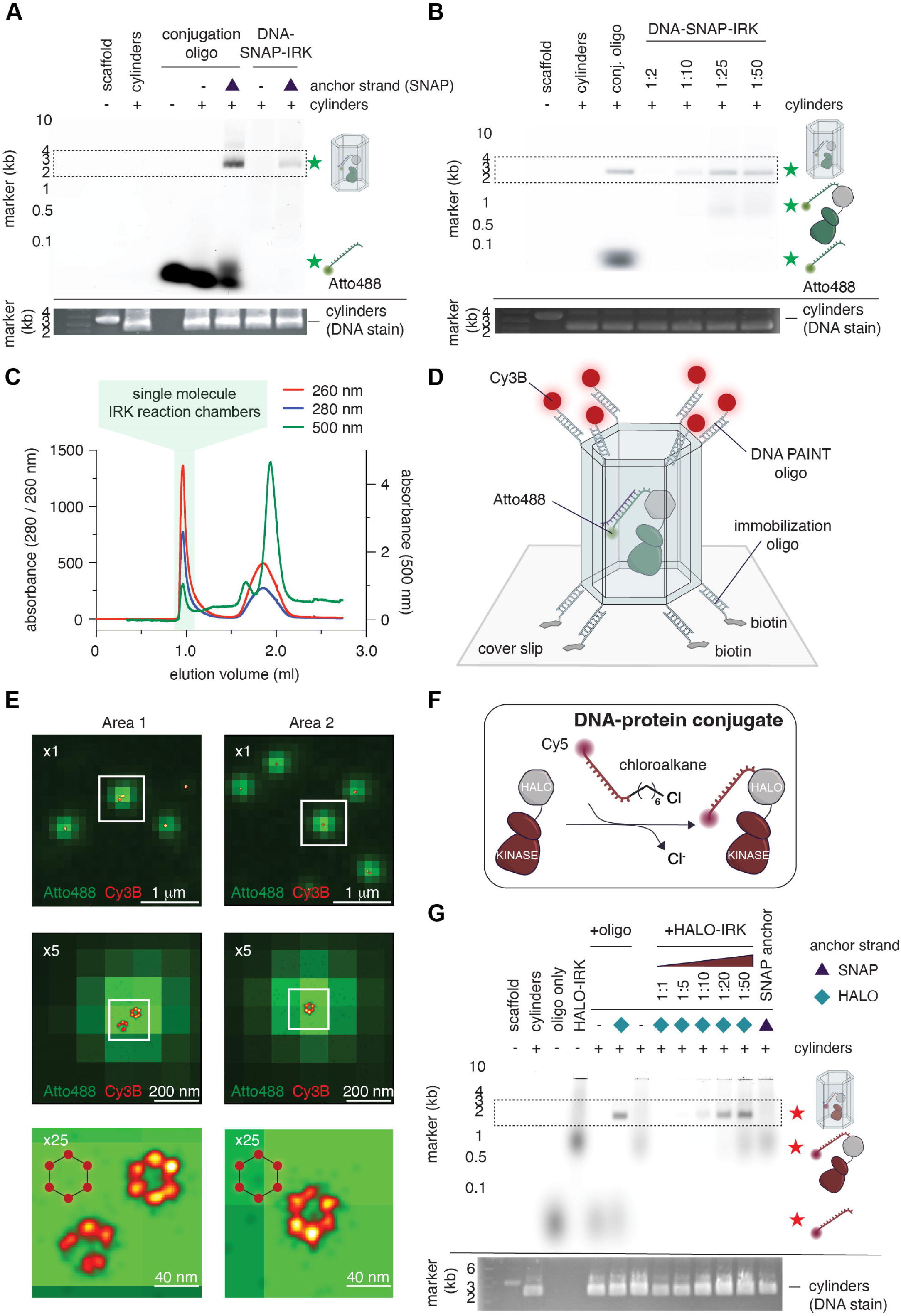
Reaction chambers can be loaded in a sequence-specific manner. A. Site-specific incorporation of the conjugation oligo into the origami cylinders dependent on the presence of the anchor strand. Incorporation of conjugated protein is less efficient, but also site-specific. Origami cylinders are detected by UV using the DNA stain SYBR-Safe. B. Titration of the DNA protein conjugate reveals that a 25-fold stochiometric excess of it over origamis is necessary for their full occupancy. C. Immobilized DNA protein conjugate inside the origami cylinders can be separated by size exclusion chromatography from freely diffusing one. D. The reaction chambers are immobilized on a streptavidin-coated cover slip by annealing single-stranded overhangs (x6) on one side of the cylinder to a biotinylated immobilization oligo. Single stranded-overhangs on the opposite end of the cylinder (x6) can be used to transiently bind DNA-PAINT oligos. E. Super-resolution microscopy of immobilized reaction chambers. Atto488 channel (green) is diffraction limited. Cy5 (red) channel imaged using DNA-PAINT. F. Alternative, orthogonal immobilization scheme. A HALO-tagged KD fusion protein (HALO-KD) is covalently conjugated with a single stranded oligo bearing a terminal chloroalkane. G. The HALO-IRK and SNAP-IRK DNA conjugates can be site-specifically immobilized via base pairing with complementary anchor strands.

To confirm the loading of the reaction chambers visually, we immobilized them in an upright orientation on a streptavidin-coated glass cover slip using biotinylated oligonucleotides complementary to single-strand overhangs at one end of the cylinder (Figure 2D). By employing super-resolution DNA-PAINT microscopy^33,34^, we validated the architecture of the cylinders using short, fluorescently-labeled oligonucleotides complementary to single-strand overhangs at each of the six vertices on the top of the cylinder (Figure 2D). We recovered a characteristic hexagonal pattern of fluorophores in the red channel (Cy3B) with expected dimensions (Figure 2E) and coincident with a diffraction-limited signal in the green channel (Atto488), confirming the incorporation of the conjugation strand into the cylinder (Figure 2E).

To extend the versatility of the system, we developed an immobilization strategy with which a second molecule could be site-specifically immobilized in cylinders containing a second, orthogonal anchor strand sequence (Figure 2F). Briefly, Halo-tagged IRK was covalently conjugated to an oligonucleotide bearing a 3’ Cy5 fluorophore, allowing it to be distinguished spectroscopically from reaction chambers containing Atto488-labeled SNAP-IRK conjugates. The mass difference of the tags additionally enables the separation of their respective fusion proteins by SDS-PAGE (Supplementary Figure 5A). We confirmed that SNAP-IRK and Halo-IRK DNA conjugates could be immobilized exclusively and site-specifically in their respective reaction chambers (Figure 2G).

### IRK undergoes intramolecular autophosphorylation

We next subjected single-molecule reaction chambers containing IRK to radiometric kinase assays. The cylinders were perfused with radiolabeled ATP and Mg^2+^ to initiate the reaction (Figure 3A). We observed a robust increase in autophosphorylation over time to a plateau (Figure 3B-C), demonstrating that IRK can phosphorylate itself in an intramolecular reaction. Addition of 6 μM freely diffusible kinase-dead IRK (IRK^KD^) to the cylinders resulted in a modest stimulation of intramolecular autophosphorylation of the immobilized, SNAP-IRK^WT^, as well as the accumulation of phosphorylated IRK^KD^ (Figure 3B). This indicates that the intermolecular reaction is possible at high concentrations. To rule out the possibility that the autophosphorylation of IRK immobilized inside a cylinder is the consequence of its dissociation and subsequent *trans* (intermolecular) interaction with IRK in another cylinder, we immobilized IRK^KD^, site specifically, in an orthogonal single-molecule reaction chamber using the Halo-tag strategy described previously (Figure 3F-G). Mixing of stoichiometric amounts of wild-type and kinase-dead IRK reaction chambers resulted in the exclusive phosphorylation of IRK^WT^ (Supplementary Figure 5B) with identical kinetics as IRK^WT^ chambers alone (Supplementary Figure 5C), confirming that the reaction occurs exclusively inside the chamber and, therefore, intramolecularly.

**Figure 3.**
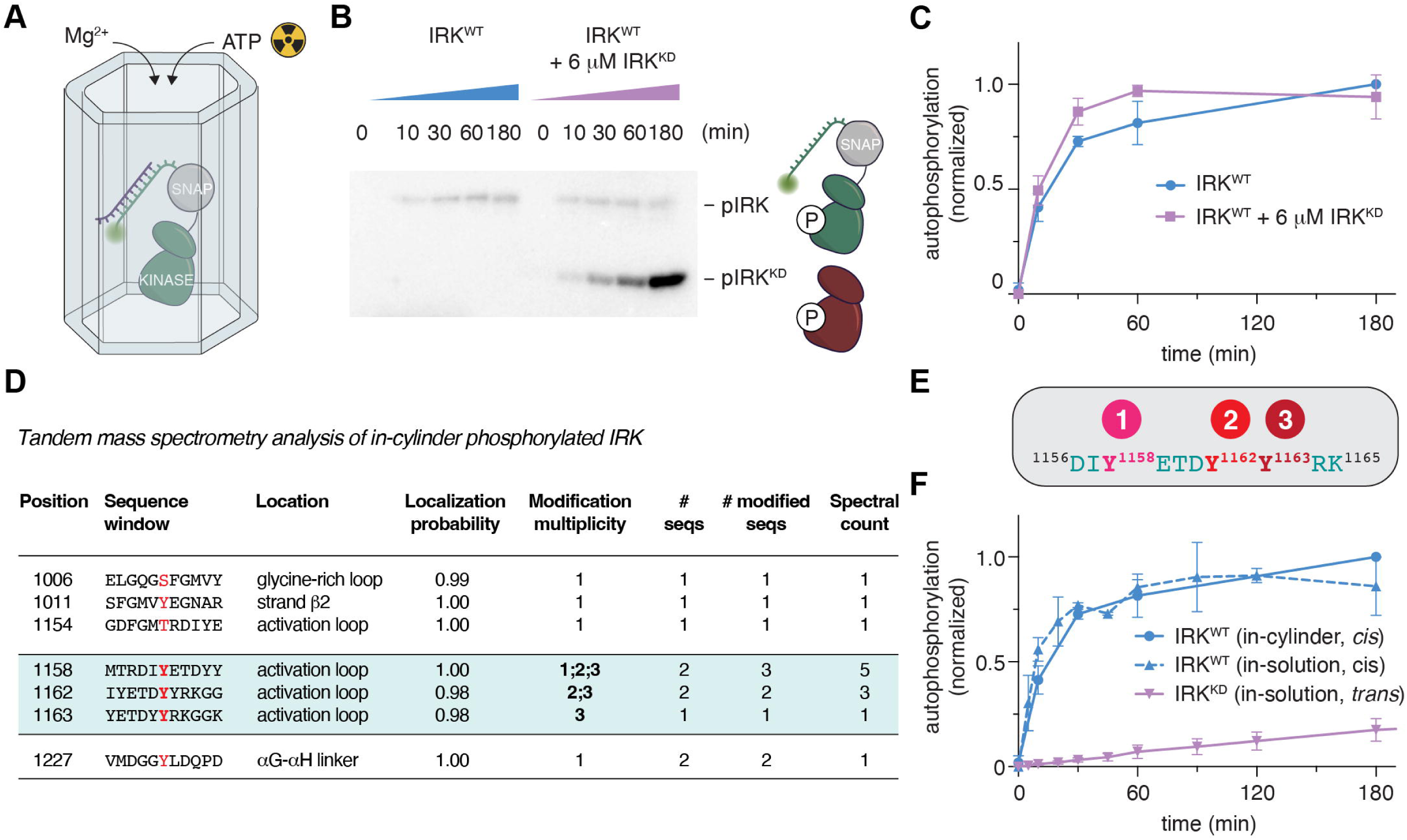
Insulin receptor kinase autophosphorylates in *cis*. A. IRK-loaded reaction chambers are perfused with radiolabeled ATP and magnesium ions to initiate the kinase reaction. B. Autoradiograph of the in cylinder single molecule reaction chamber IRK kinase assay showing increasing IRK cis-autophosphorylation over time in the presence or absence of 6 μm kinase dead IRK kinase domain (IRK^KD^). C. The presence of 6 μm IRK^KD^ stimulates autophosphorylation of active IRK. D. All three IRK activation loop tyrosines undergo autophosphorylation inside the reaction chambers. E. Based on modification multiplicity, activation loop autophosphorylation is sequential: phosphorylation of Y1158 is followed by Y1162 and Y1163. F. In-cylinder autophosphorylation kinetics recapitulate the kinetics of freely diffusing protein, while the intermolecular (*trans*) reaction is markedly slower.

To identify the residues in IRK undergoing *cis*-autophosphorylation, we subjected the IRK recovered from the in-cylinder kinase assay to tandem mass spectrometry. We detected robust phosphorylation of all three canonical tyrosines (Y1158, Y1162 and Y1163) as well as the relatively weak off-target phosphorylation of residues proximal to the catalytic center of the kinase (Figure 3D). Moreover, since we sampled a time point before the reaction reached completion and all three tyrosines reside on the same tryptic peptide, we used the stoichiometry and phosphorylation site identification (modification multiplicity) of these peptides to infer the order of phosphorylation events. This analysis revealed a sequential mechanism in which Y1158 is phosphorylated first, followed by Y1162 and, finally, Y1163 (Figure 3D-E). This is consistent with our MD simulations, in which Y1158 consistently exhibited the most persistent ATPγP-proximal sampling (Supplementary Figure 1H).

We next compared the reaction kinetics of in-cylinder IRK autophosphorylation with those of freely diffusing IRK. The kinetics were indistinguishable, demonstrating that the reaction chamber itself does not affect the intrinsic kinetics of the reaction and that bulk solution kinetics predominantly reflect intramolecular autophosphorylation (Figure 3F). In contrast, the intermolecular, *trans* reaction, normalized by protein concentration, exhibited linear kinetics with a significantly slower rate.

Finally, to understand how IRK autophosphorylation is controlled allosterically, we examined the role of the juxtamembrane (JM) segment, a well-established regulatory element^14,35^. Deletion of the JM segment (IRK^ΔJM^) completely abolished autophosphorylation (Supplementary Figure 5D-E), demonstrating that this region is essential for activation loop autophosphorylation. In summary, activation loop autophosphorylation of IRK is intrinsically an intramolecular reaction (*cis*) and it depends, allosterically, on a membrane proximal segment that links it, via a transmembrane helix, to the insulin-sensing ectodomains of the insulin receptor.

### Allosteric activation of intramolecular autophosphorylation

Historically, concentration-dependence of protein kinase autophosphorylation has been widely used to distinguish between intra- and inter-molecular autophosphorylation^18–20^. A concentration-dependent increase in autophosphorylation kinetics is conventionally interpreted as evidence for an intermolecular *trans*-reaction as opposed to the pseudo-first order intramolecular *cis* reaction, which logically appears as if it should be intrinsically concentration independent. This interpretation, however, has one caveat: it cannot discriminate intermolecular phospho-transfer from a potential allosteric effect of dimerization that promotes intramolecular *cis*-autophosphorylation. The same concentration dependence would be expected in both cases.

To demonstrate this effect explicitly, we turned to the bacterial kinase PknB of *Mycobacterium tuberculosis*. We chose PknB for multiple reasons: (1) structurally, its kinase domain has been shown to form a back-to-back dimer that is incompatible with a *trans* reaction within a dimer^36^ (Figure 4A), (2) residues in the back-to-back interface have been shown to influence kinase activity^37^, and (3) it belongs to an evolutionarily ancient branch of the protein kinase family^3,38^. *Cis* autophosphorylation of PknB would suggest that intramolecular activation loop autophosphorylation may be more broadly conserved than previously appreciated. PknB exhibited a concentration-dependent increase in autophosphorylation in bulk solution (Figure 4B), a result that has previously been interpreted as evidence for an intermolecular reaction^37,39^. To determine the nature of the concentration-dependent effect, we performed kinase assays in which the concentration of catalytically active SNAP-tagged PknB was kept constant while increasing the concentration of kinase dead, untagged PknB in the reaction (Supplementary Figure 6A). After separation of the two proteins by SDS-PAGE, we could quantify phosphorylation of the two proteins independently (Supplementary Figure 6B-E). Increasing concentrations of kinase-dead PknB stimulated autophosphorylation of the wild-type kinase until a concentration at which further stimulation of the reaction was not possible (Figure 4C). Mutation of a single residue in the back-to-back dimer interface of PknB (PknB^KD^ ^L33D^) was sufficient to completely abolish this stimulatory effect (Figure 4C, Supplementary Figure 6B-E). We confirmed that autophosphorylation of PknB^WT^ resulted predominantly in phosphorylation of a single residue (Supplementary Figure 6F), which we confirmed to be either the canonical activation loop residue, T173, or the neighboring T171 (Figure 4D, Supplementary Figure 6G). Autophosphorylation gave rise to negligible amounts of di-phosphorylated PknB^WT^ (Supplementary Figure 6F), indicating that phosphorylation of T171 and T173 are mutually exclusive. Importantly, we observed that kinase-dead PknB could be phosphorylated under the conditions of this assay (Supplementary Figure 6B-D), demonstrating that the *trans* reaction, as previously observed for IRK, is possible at high protein concentrations. However, PknB^KD^ bearing the L33D mutation exhibited a considerably lower rate of intermolecular *trans*-phosphorylation compared to PknB^KD^ (Supplementary Figure 6E), indicating that the rate of the intermolecular reaction depends critically on the prior allosteric activation of wild-type PknB. In summary, neither a concentration-dependent increase in activation loop autophosphorylation nor observation of *trans*-autophosphorylation in bulk solution can be taken as unambiguous evidence of a physiologically relevant intermolecular reaction.

**Figure 4.**
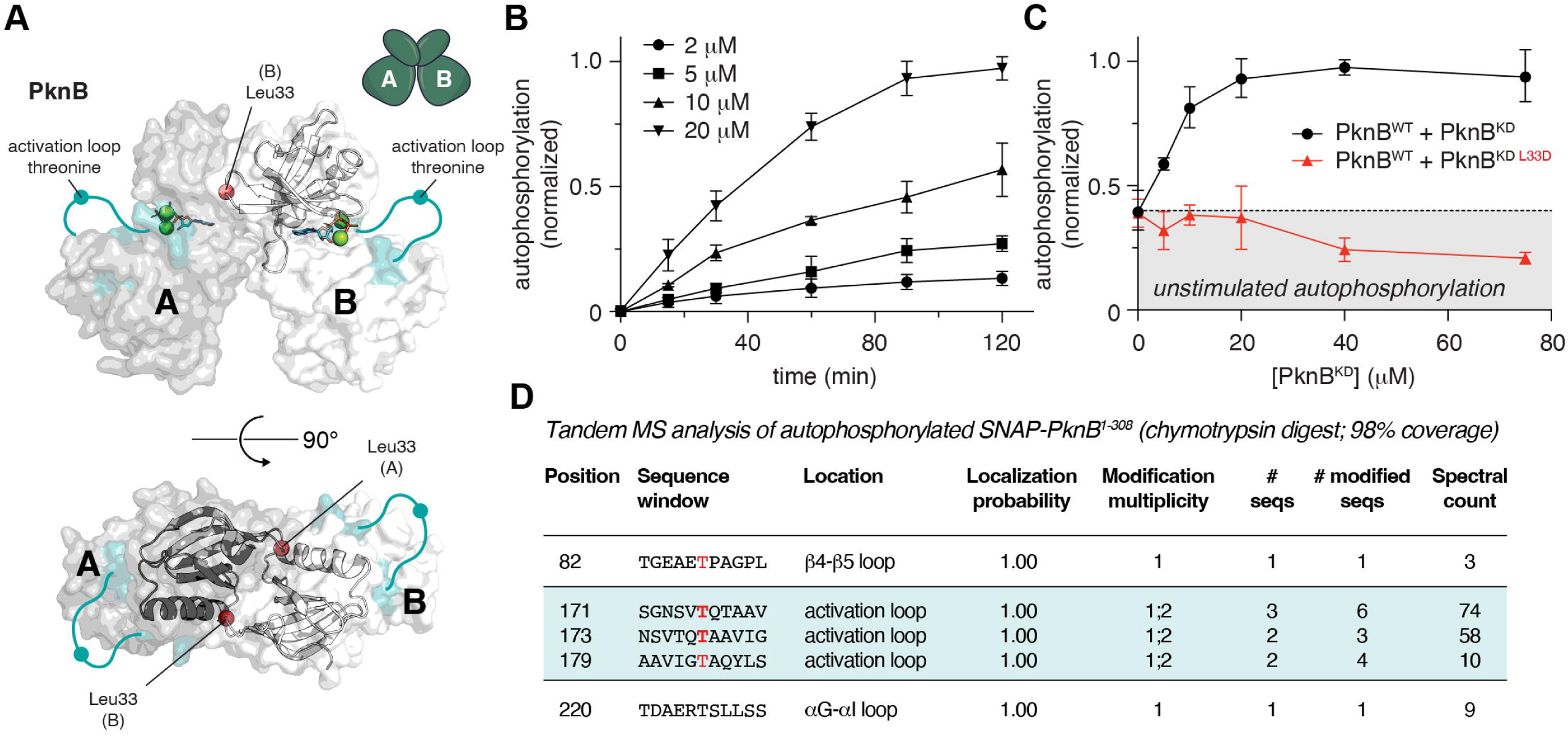
Allosteric activation of intramolecular autophosphorylation. A. Crystal structure of *M. tuberculosis* PknB (1MRU)^36^. PknB crystallized as a dimer with two molecules per asymmetric unit. The dimer interface is centered on the N-lobe of the kinase domain and the invariant Leu33 (red sphere) which inserts its side chain into a hydrophobic pocket between αC and β4/β5 in the N-lobe of the opposing protomer. B. Concentration-dependence of PknB autophosphorylation. C. Stimulation of PknB autophosphorylation by kinase-dead PknB but not by kinase-dead PknB^L33D^. D. Intact mass spectrometry analysis of the reaction product. E. Tandem mass spectrometry of the reaction product upon trypsin digest. F. Tandem mass spectrometry of the reaction product upon chymotrypsin digest.

## Discussion

To date, conventional biochemical assays and crystal structures of protein kinases have predominantly been interpreted as evidence of intermolecular activation loop autophosphorylation. Our findings challenge this notion by demonstrating that intramolecular activation loop autophosphorylation is the dominant activation mechanism of a human tyrosine kinase and that intramolecular autophosphorylation can be stimulated, allosterically, by dimerization in a bacterial serine/threonine kinase.

Fundamentally, this work highlights the limitations of the experimental approaches that have historically been used to infer *trans*-autophosphorylation. The detection of *trans*-phosphorylation of a kinase-dead mutant, particularly under conditions of high protein concentration, demonstrates only that intermolecular phospho-transfer is possible. This is, however, not surprising, because the enzyme, substrate (activation loop), and reaction mechanism are all chemically identical, irrespective of whether the reaction occurs in *cis* or *trans*. A kinase capable of *cis*-autophosphorylation would therefore be expected to be able to do so in *trans* as long as the activation loop is sufficiently accessible and the molecules are free to interact. Thus, the ability of a kinase to catalyze *trans*-autophosphorylation is not evidence of its inability to do so in *cis*.

Likewise, a concentration-dependent increase in kinase activity is evidence of dimerization-dependent activation, not evidence of *trans*-autophosphorylation. This is exemplified by kinases that depend on allosteric activation via back-to-back dimerization^25,40,41^. In the case of PknB, autophosphorylation is allosterically stimulated by dimerization in a back-to-back configuration that is incompatible with an intermolecular, *trans* reaction within the same dimer. Dimerization stabilizes the active conformation, thereby facilitating phospho-transfer, intramolecularly, within each protomer. Consistent with intramolecular autophosphorylation, previous MD simulations of PknB detected only *cis* interactions over the time course of the simulation^42^. The kinase domains of the four mammalian ISR kinases exhibit back-to-back dimerization that is structurally homologous to PknB^25,40,43,44^ and have recently been shown to be activated by dimerization-dependent *cis*-autophosphorylation^25^. Together, these observations illustrate that intermolecular regulation of kinase activity is not the same as intermolecular phospho-transfer.

Numerous crystal structures have captured activation loop-exchanged kinase domain dimers^5,10^. These assemblies have been interpreted as snapshots of *trans*-autophosphorylation^17^. However, they might also reflect thermodynamically favorable crystal-packing arrangements^5,45^. Our simulations underscore why static structures alone cannot unambiguously distinguish between intra- and inter-molecular autophosphorylation by revealing that IRK can still undergo the minimal conformational changes required for *cis*-autophosphorylation. Curiously, although the field has since coalesced around the intermolecular model of IRK activation, an early study found the activation mechanism to be intramolecular in nature^46^. Although structures of the full-length IR have been determined by cryo-electron microscopy, none have captured the kinase domains at high resolution^47,48^. More sophisticated reconstitution experiments, possibly in complex with adaptor proteins such as IRS1/2^49,50^, are likely required to resolve how insulin binding is coupled to activation loop autophosphorylation.

Our findings raise the question of how widely intramolecular activation loop autophosphorylation is conserved across the kinome, given that both IRK and PknB most likely share a common ancestor^1,3^. Whilst we cannot exclude the possibility that some kinases may have evolved an activation mechanism that depends on intermolecular *trans*-autophosphorylation of their activation loops, logic favors an intramolecular reaction for the following reasons: (1) intermolecular *trans*-autophosphorylation would require a mechanism to prevent intramolecular *cis*-autophosphorylation while favoring intermolecular phosphorylation. The necessary length and flexibility of the activation loop that would be required for it to be engaged by the catalytic center of a neighboring protomer, however, would equally likely enable an intramolecular reaction. Since the substrate phosphorylation site is identical in both cases, effective local concentration dictates that, if sterically feasible, the intramolecular reaction will dominate; (2) intramolecular autophosphorylation would, hypothetically, require only an allosteric input, as opposed to the diffusion, encounter, and productive interaction of two copies of the same protein in the crowded cellular environment ^51,52^. Given that protein kinases are generally expressed at low nanomolar concentrations^53^, unless they exist as pre-formed dimers, mathematical modeling of their encounter kinetics suggests that acute and high-fidelity signal transduction in a cell is not possible^7^. This principle applies equally to intermolecular *trans*-autophosphorylation and dimerization-dependent intramolecular *cis*-autophosphorylation. In case of the sequential activation of kinases in a cascade, it implies that they must be scaffolded together^54^, irrespective of whether activation loop phosphorylation occurs in *trans* (between kinases) or, allosterically, in *cis*; (3) an intermolecular reaction necessarily requires resolution of a face-to-face dimer prior to substrate engagement, a step which is not required for a kinase that undergoes intramolecular activation loop autophosphorylation (Figure 5). The back-to-back conformations of PknB, the ISR and UPR kinases, and Nek7/Nek9 are simultaneously compatible with dimerization-mediated allosteric stimulation of both activation loop and downstream substrate phosphorylation^36,40,55^. Consistent with auto- and substrate phosphorylation being allosterically coupled, the same allosteric input required for intramolecular activation loop autophosphorylation of PKR-like endoplasmic reticulum kinase (PERK) is required for subsequent phosphorylation of its substrate, eIF2α^25^. If activation loop autophosphorylation is intramolecular by nature, it is, by definition, mechanistically distinct from intermolecular substrate phosphorylation. As such, activation loop autophosphorylation must precede substrate phosphorylation, thereby resolving the enigma of autophosphorylation (Figure 5).

**Figure 5.**
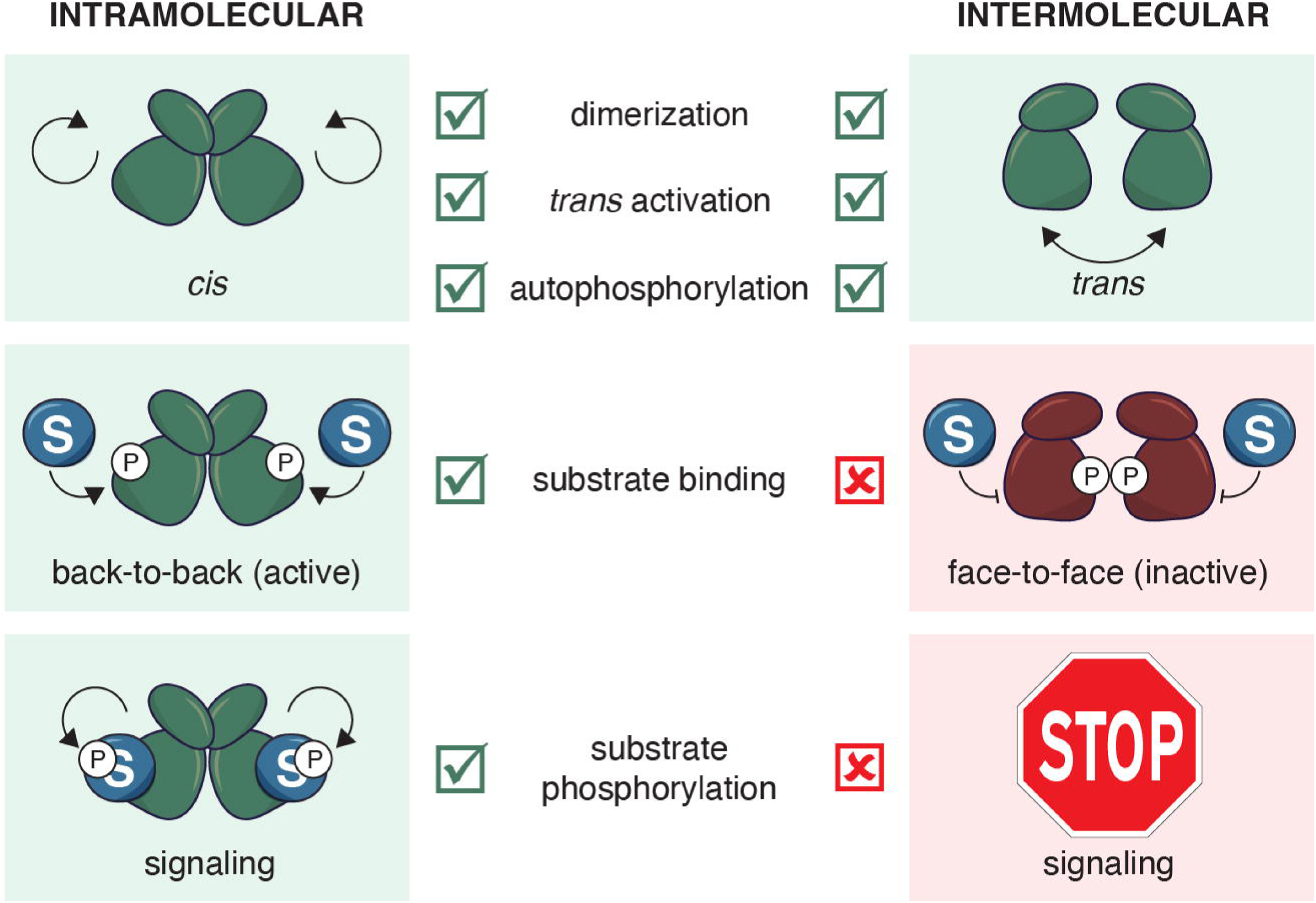
Intra- vs inter-molecular protein kinase activation. Comparison of intra-(left) and inter- (right) molecular protein kinase activation by autophosphorylation. Intramolecular (*cis*) autophosphorylation can be allosterically stimulated in *trans* and is compatible with immediate immobilization of the phosphorylated activation loop, consequent substrate binding, and substrate phosphorylation. In contrast, intermolecular (*trans*) autophosphorylation is incompatible with substrate binding or phosphorylation without an intermediate step to resolve the face-to-face dimer required for activation loop phosphorylation.

Beyond resolving a longstanding question in kinase biology, our work establishes a DNA origami platform for distinguishing between inter- and intra-molecular enzymatic reactions at the single-molecule level. The modularity of this platform makes it adaptable to other systems, such as E3 ubiquitin ligases, for which distinguishing between inter- and intra-molecular catalysis remains an open mechanistic question in many cases^56,57^. Moreover, the use of orthogonal anchor strand sequences enables the site-specific positioning of multiple proteins within the same reaction chamber. This toolbox expands the range of mechanistic questions that can be addressed by enabling one to distinguish regulatory interactions from catalytic events, which is not possible in conventional bulk biochemical assays.

## Methods

### DNA origami design

DNA origami reaction chambers were designed using caDNAno 0.2.3^58^ in a honeycomb lattice from 72 helices that form a hexagonal cross-section. Dimensions of the reaction chamber were chosen such that they allow enough space for a DNA-protein kinase conjugate to be immobilized inside the cylinder. Attachment sites were placed in the middle of the cylinder to ensure immobilized proteins could not reach outside of the cylinder. Staples were broken manually using the following rules: 1) staple length must be between 35 and 60 bases; 2) each staple strand must contribute to at least one continuous double-stranded segment of 10 bp or longer; 3) if possible, a staple strand should not feature one or more short (7bp) segments between two longer segments; 4) if possible, segments shorter than 7 bp should be avoided; 5) never break both strands at a crossover. 3D models and all-atom structures were generated using the CanDo webserver^59^.

### Fabrication of DNA cylinders

Reaction mixtures contained 200 nM staples (IDT), 20 nM 8064 scaffold (tilibit), 15 mM MgCl_2_, 5 mM Tris pH 8, 5 mM NaCl and 1 mM EDTA. The sample was heated for 15 min to 65°C before cooling it down at 1°C per hour from 60 °C to 40 °C. To remove excess staple strands, the origamis were washed in a 100 kDa cutoff concentrator (Merck) five times with wash buffer (5 mM NaCl, 5 mM Tris pH 8, 8 mM MgCl_2_) before being transferred into storage buffer (50 mM HEPES pH 7.4, 150 mM NaCl, 10 mM MgCl_2_) in DNA LoBind tubes (Eppendorf). Successful assembly of origamis and removal of excess staples was evaluated by agarose gel electrophoresis (2% agarose, 1x TAE, 10 mM MgCl_2_) with a running buffer of 1x TAE supplemented with 10 mM MgCl_2_ at 80 V for 90 min.

### Negative stain electron microscopy

400-mesh copper grids (Quantifoil) coated with a thin carbon film (homemade) were glow-discharged for 60 s using a Bal-Tec SCD 005. A 5 μL aliquot of cylinders (100 nM in 5 mM MgCl_2_, 5 mM Tris pH 8 and 5 mM NaCl) was applied to the grid and incubated for 60 s. The grid was blotted with Whatman filter paper and immediately stained with 5 μL of 2.0 % (w/v) uranyl acetate in water for 60 s. The staining step was repeated once. The grids were blotted dry and stored in a vacuum chamber until inspection. Images were recorded at a nominal magnification of 71,000x (0.856 nm/pixel) using a FEI Morgagni 268D microscope operated at 80 kV, equipped with an Olympus-SIS MegaView III camera.

### Electrophoretic mobility shift assay (EMSA)

The stoichiometric loading of the origami cylinders was determined by electromobility shift assay (EMSA). The indicated ratios of either conjugation oligo or conjugated protein and DNA origamis were mixed and incubated overnight at room temperature. EMSAs were run on agarose gels (2% agarose, 1x TAE, 10 mM MgCl_2_) at 80 V for 90 min in a water bath to avoid heating of the gel. Atto488 or Cy5 fluorescence was imaged with an Amersham Typhoon imager before subjecting the gel to SYBR safe stain (10 uL SYBR Safe in 15 mL TAE) for 25 min shaking at room temperature and imaging with a Fusion FX7 imager (Vilber Lourmat). The fluorescence signal was quantified in Image J.

### Radiometric kinase assays

Origamis were incubated with a 25x molar excess of the SNAP fusion protein or a 50x molar excess of the Halo fusion protein overnight at room temperature. To remove the unincorporated protein, the sample was applied onto a S6 3.2/300 size exclusion column (Aekta Micro system) equilibrated in kinase assay buffer (50 mM HEPES pH 7.4, 150 mM NaCl, 10 mM MgCl_2_, 1 mM TCEP). Protein-derivatized cylinders were pooled and concentrated to 100 nM using 100 kDa cutoff concentrators (Merck Millipore).

Radiometric kinase assays were performed using radiolabeled [γ^32^ P] ATP (Hartman Analytic) at room temperature with a final ATP concentration of 1 mM, spiked with 1:20 [γ^32^ P] ATP. All reactions contained 2 mM MgCl_2_, with the exception of PknB assays, in which MgCl_2_ was substituted with MnCl_2_. For all assays that contained more than one protein, the proteins were mixed and preincubated for 1 h at room temperature before starting the assay. At each time point, 10 uL of sample was collected, quenched with 2.5 μL of 1 M EDTA pH 8 and 3 μL 5x SDS loading dye, and immediately boiled at 95 °C for 5 min.

### Super-resolution microscopy sample preparation

40 μL of BSA-biotin (5 mg/mL in PBS; Sigma) were spotted on Parafilm and a freshly plasma-cleaned cover slip was incubated face down for 15 min at room temperature. The cover slip was then rinsed with PBS and incubated with 40 μL Neutravidin (10 mg/mL in PBS; ThermoFisher before being rinsed again with PBS. 40 μL of 750 pM DNA origami in PBS were applied to the parafilm and the coverslip was incubated for 15 min before being washed in PBS and incubated once again with 40 μL BSA-biotin (5 mg/mL in PBS) for 5 min to block unspecific binding of Cy3B-labeled DNA imager strands to the sample. The glass support with the immobilized origami was then mounted on a custom magnetic metal plate with a hole in its center where the coverslip is stably held in place by a ring magnet encased in a PVC shell. An O-ring mounted in the outer shell of the magnet ensures the liquid tightness of the construction, such that imager strand-containing buffer can be placed on the sample. Imaging buffer (Massive Photonics) with the addition of imager strands (1nM final concentration) was then applied to the sample.

Prior to sample preparation for microscopy, the DNA origamis were incubated overnight with a 2.5-fold excess of Atto488-labelled conjugation oligo and 1 hour before sample preparation with a 12-fold excess of biotinylated oligo for immobilization (two-fold stoichiometric excess over the six overhangs per cylinder). Further details on the imaging and reconstruction of DNA-PAINT images are reported in the Supplementary Material.

## Acknowledgements

Proteomics analyses were performed on instruments of the Vienna BioCenter Core Facilities (VBCF). Negative stain electron microscopy was supported by the Electron Microscopy Facility of the Vienna BioCenter Core Facilities (VBCF). Cell culture medium was prepared by the Max Perutz labs media kitchen. We thank Dr. W. Todd Miller for critical feedback on the manuscript. This work was funded by an Out-of-the-Box Award of the Vienna Biocenter PhD Program to E.K. and Austrian Science Fund (FWF) grants P36212 and P36724 to T.A.L. and P30550 to BZ.

## Author contributions

E.K. expressed, purified and characterized all proteins employed in this study with help from A.E. C.S. performed all MD simulations and analyzed the data together with B.Z. F.R. collected all super-resolution imaging data with the help of E.K. and analyzed the data together with J.R. F.P. designed the DNA origami cylinder. L.P. collected electron microscopy images of the origami cylinders. B.Z., F.P., and T.A.L. obtained funding for the work. E.K. wrote the manuscript, which was edited by all authors.

## Data availability

The mass spectrometry proteomics data have been deposited to the ProteomeXchange Consortium via the PRIDE^60^ partner repository with the dataset identifiers PXD083365 and PXD083367.

