## Supplementary Information for "Single-molecule reaction chambers reveal intramolecular protein kinase activation"

Contents:

Supplementary Figures 1-6

Supplementary Methods

Supplementary References

### Supplementary Figures

#### Supplementary Figure 1. Protein kinase activation loop *cis*-autophosphorylation is sterically possible.

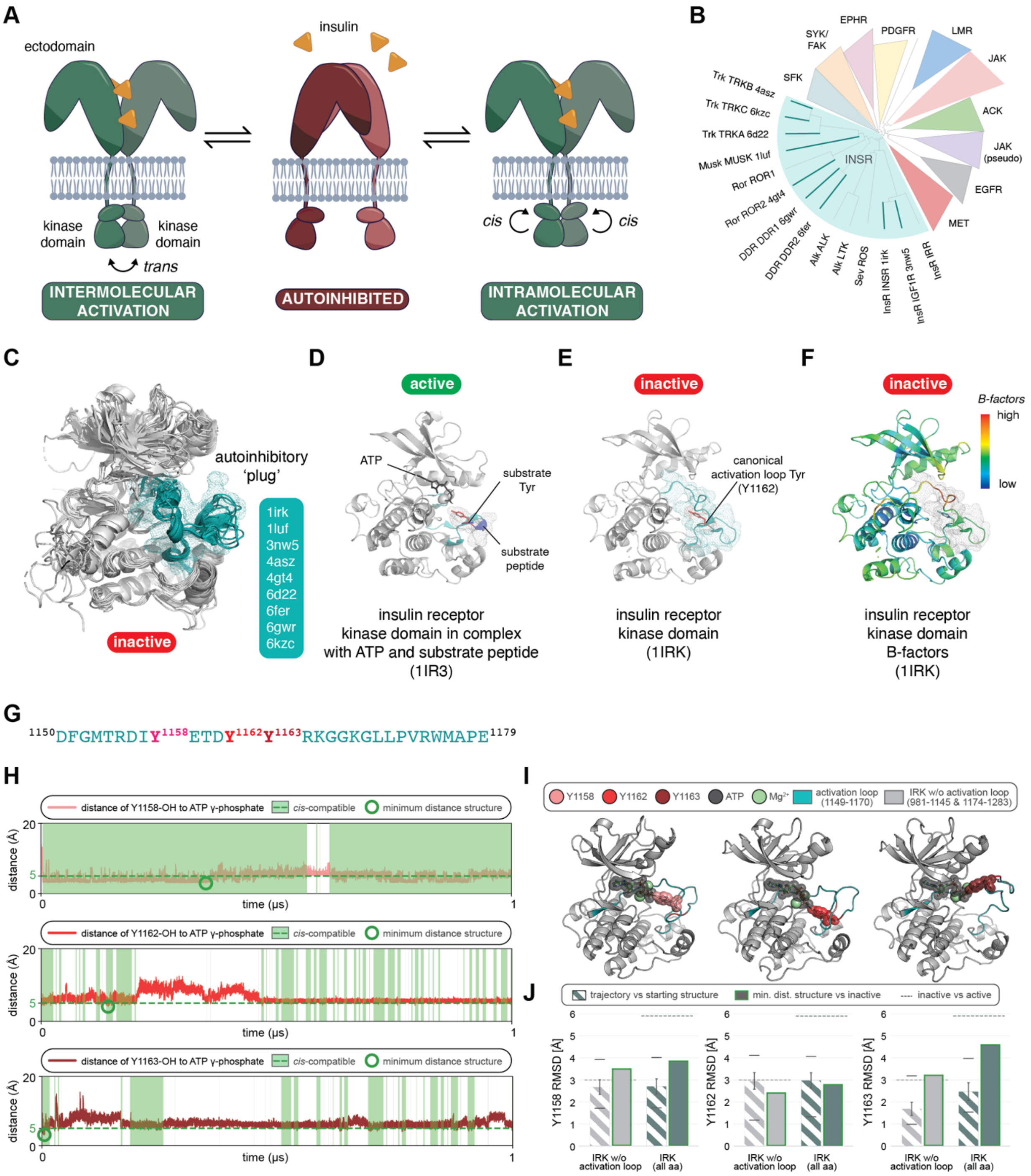

A. Plausible models for activation of the insulin receptor (IR) by *cis*- or *trans*-autophosphorylation.
B. Phylogenetic tree of the receptor tyrosine kinases.
C. Experimental structures of the inactive conformation of the kinase domain of the IR and related kinases indicated with the bold lines in panel B. The activation loop is depicted in teal mesh.
D. Crystal structure of IRK in complex with ATP and substrate peptide (PDB:1IR3).
E. Crystal structure of apo-IRK without any ligands (PDB:1IRK).
F. B-factors of the apo-IRK crystal structure (PDB:1IRK) from panel E.
G. Peptide sequence of the IRK activation loop that contains the three canonical autophosphorylation sites.
H. Molecular dynamics simulations of *cis*-autophosphorylation-competent IRK conformations. Distance between the hydroxyl oxygen of each phosphate-acceptor tyrosine and the ATP gamma phosphate phosphorus atom (ATPγP) over fully unrestrained 1 μs simulations (Y1158 top, Y1162 middle, Y1163 bottom). Dashed green: 5 Å threshold for *cis*-autophosphorylation compatibility; green bands: sub-threshold intervals; open green circle: minimum-distance frame.
I. Minimum-distance conformations of IRK with ATP-$Mg^{2+}$ in the ATP-binding pocket with Tyr-OH to ATPγP distances of 2.96 / 4.08 / 3.14 Å for Y1158 / Y1162 / Y1163, respectively.
J. Backbone RMSD for IRK without and with the activation loop included. Striped bars: final 90% of trajectory vs. first frame (mean ± s.d., min/max markers). Green-bordered bars: minimum-distance structure vs. inactive IRK[9]. Dashed line reference: RMSD between inactive[9] (PDB:1IRK) and active phosphorylated (PDB:1IR3)[27] IRK crystal structures.

### Supplementary Figure 2. Fabrication of single-molecule reaction chambers from DNA origami.

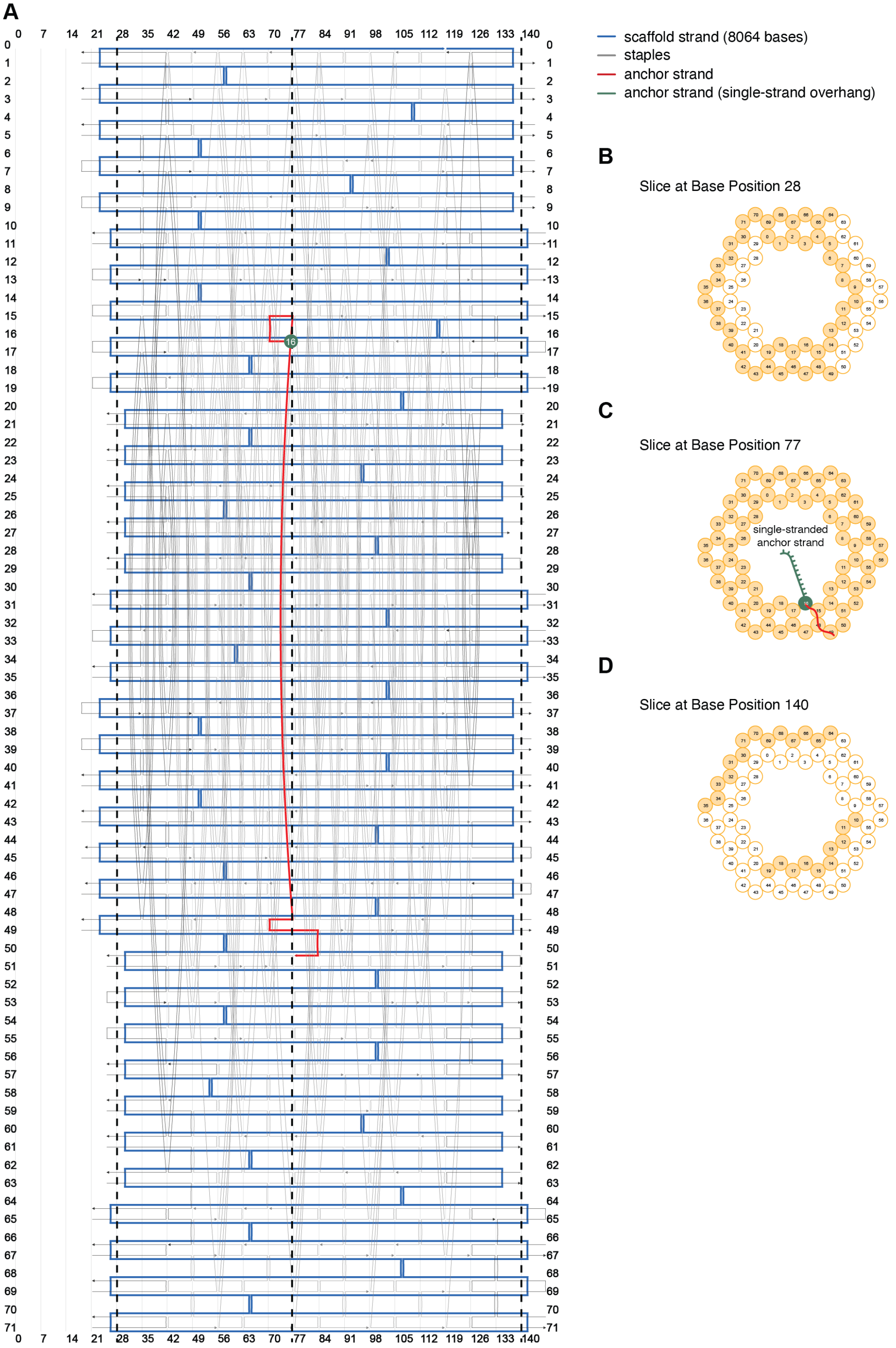

A. CaDNAno model of the origami cylinders employed in this study. Scaffold strand is indicated in blue, staple strands in grey, the staple part of the anchor strand in red, and the start of its single-stranded overhang in green.
B. Cross-section of the origami cylinder at nucleotide position 28. Complete double helices are indicated by filled circles, absent helices by open circles.
C. Cross-section of the origami cylinder at nucleotide position 77, indicating the staple part (red) and the single-stranded overhang (green) of the anchor strand. The anchor strand is wound around helices 49, 48 and 15 before its single-stranded part reaches into the inside of the cylinder from helix 16.
D. Cross-section of the origami cylinder at nucleotide position 140. Complete double helices are indicated by filled circles, absent helices by open circles.

### Supplementary Figure 3. Mass spectrometry of proteins employed in this study.

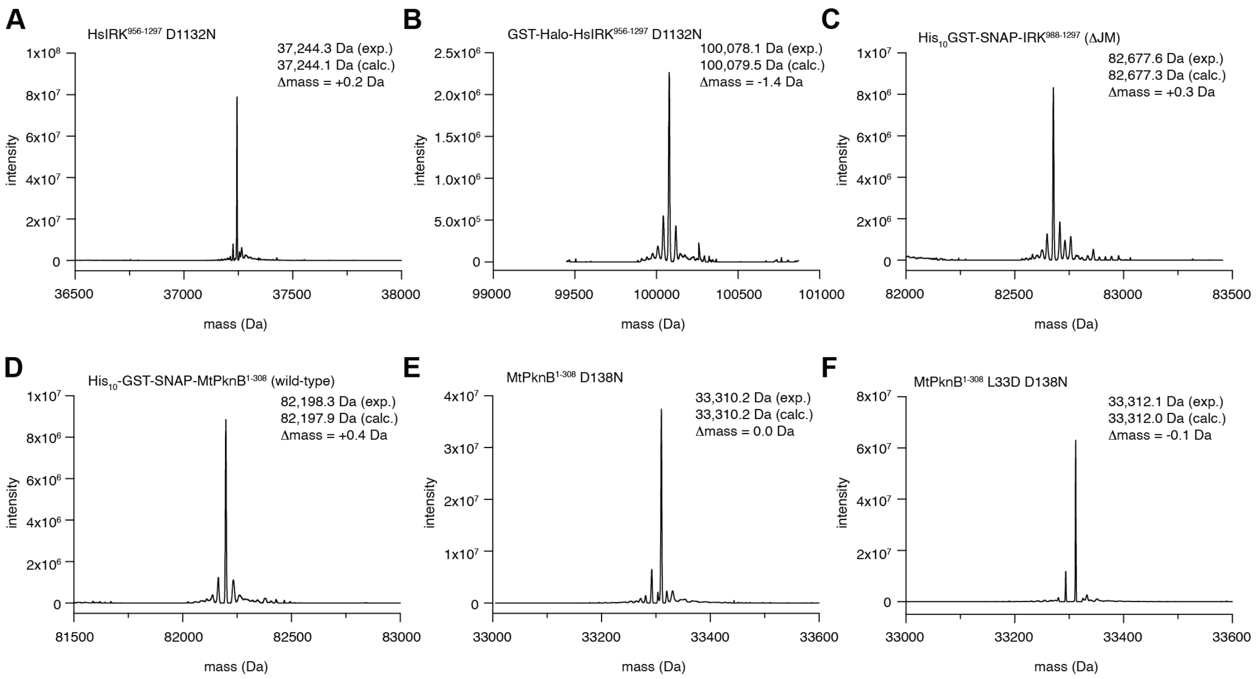


A. Intact mass spectrum of HsIRK$^{956-1297}$ D1132N.
B. Intact mass spectrum of GST-Halo-HsIRK$^{956-1297}$ D1132N.
C. Intact mass spectrum of His-GST-SNAP-HsIRK$^{988-1297}$ (ΔJM).
D. Intact mass spectrum of His-GST-SNAP-MtPknB$^{1-308}$.
E. Intact mass spectrum of MtPknB$^{1-308}$ D138N.
F. Intact mass spectrum of MtPknB$^{1-308}$ L33D D138N.

#### Supplementary Figure 4. Reaction chambers can be loaded in a site-specific manner.

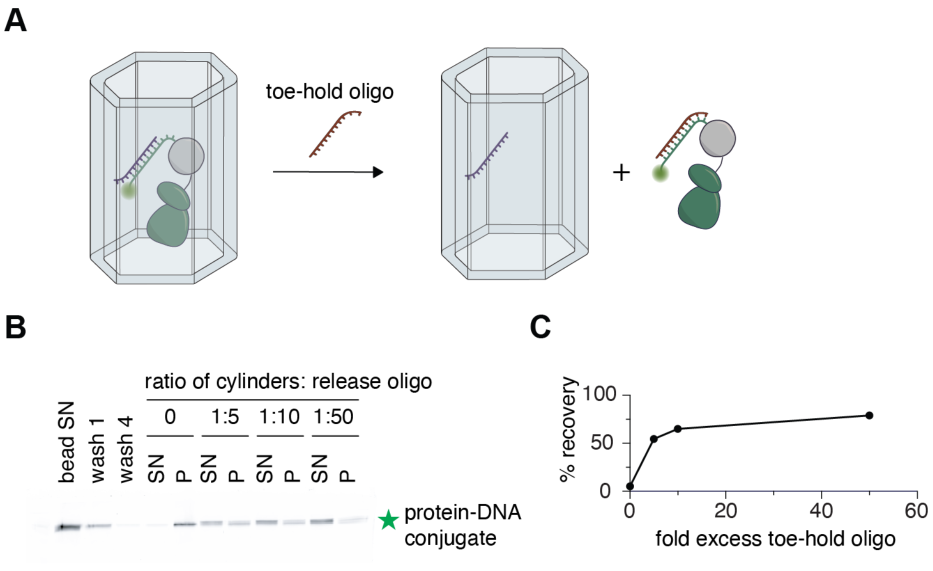


A. Immobilized DNA-protein conjugate can be competitively displaced from the chambers by the addition of a toe-hold oligo with a sequence complementary to the conjugation oligo.
B. Titration experiments with reaction chambers immobilized on beads show that upon addition of a molar excess of toe-hold oligo, the DNA-protein conjugate is eluted from the chambers.
C. Quantification of the titration experiments from panel B.

#### Supplementary Figure 5. IRK undergoes intramolecular autophosphorylation.

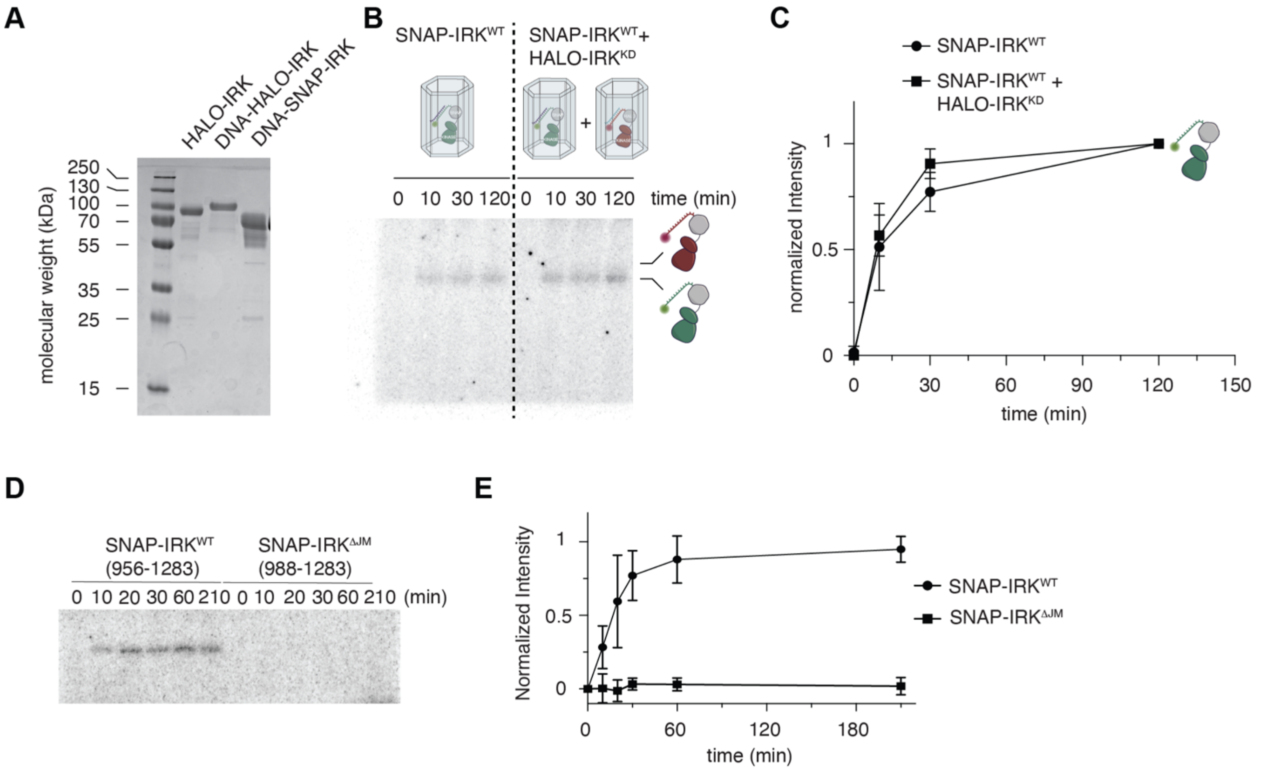


A. SDS-PAGE of DNA-HALO-IRK and DNA-SNAP-IRK conjugates.
B. Autoradiograph of IRK autophosphorylation. Left: DNA-SNAP-IRK$^{WT}$ cylinders alone. Right: stoichiometric amounts of DNA-SNAP-IRK$^{WT}$ and DNA-HALO-IRK$^{KD}$ chambers were mixed.
C. Quantification of the autoradiograph in B.
D. Autoradiograph of IRK autophosphorylation, comparing IRK$^{956\text{-}1283}$ and IRK$^{988\text{-}1297}$ (ΔJM).
E. Quantification of the autoradiograph in D.

#### Supplementary Figure 6. Allosteric activation of intramolecular autophosphorylation.

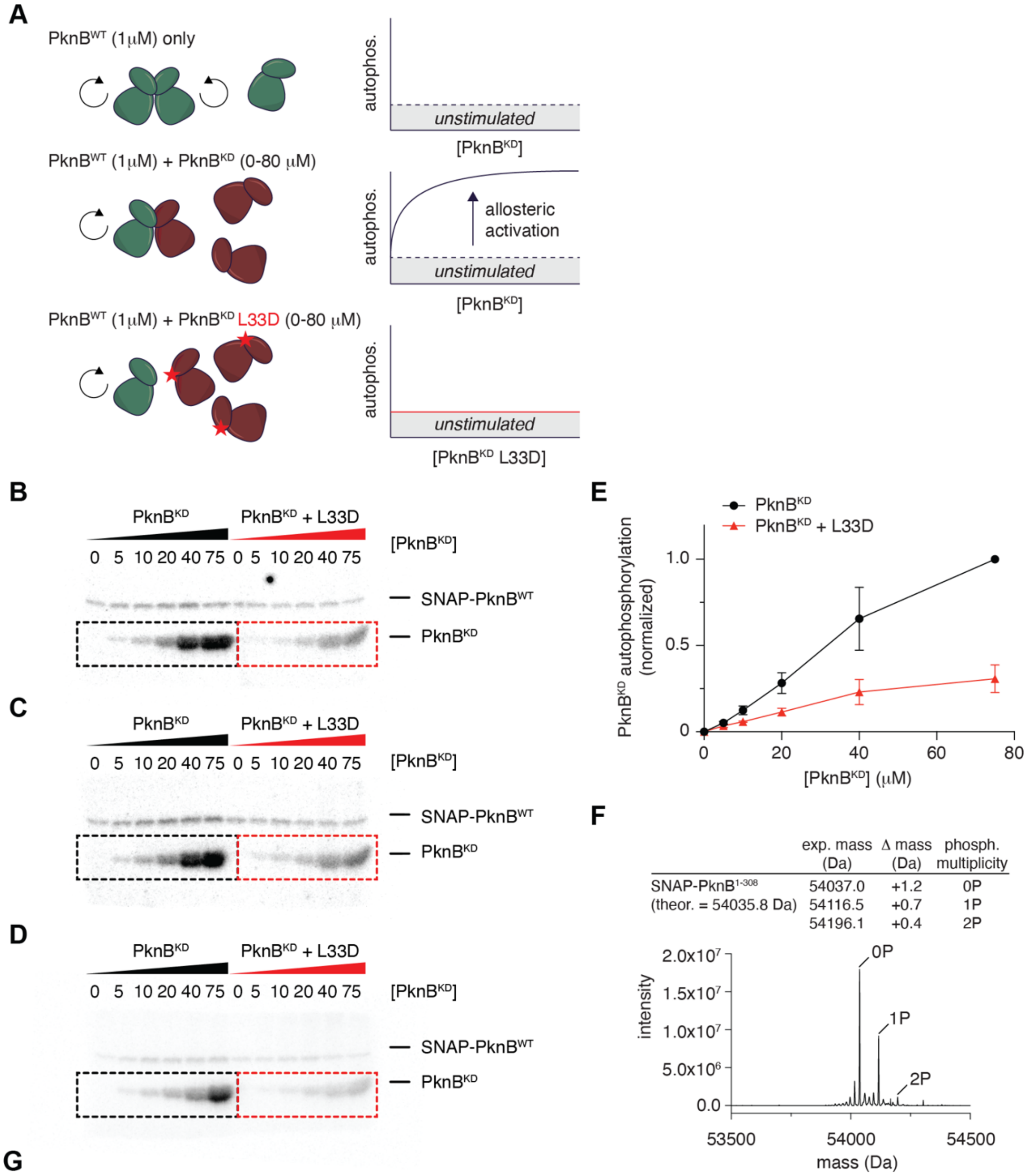


**G**

*Tandem MS analysis of autophosphorylated SNAP-PknB$^{1-308}$ (trypsin digest; 72% coverage)*

| Position | Sequence window | Location | Localization probability | Modification multiplicity | # seqs | # modified seqs | Spectral count |
|---|---|---|---|---|---|---|---|
| 149 | IMISATNAVKV | catalytic loop | 1.00 | 1 | 1 | 2 | 17 |
| 169 | ADSGNSVTQTA | activation loop | 1.00 | 1;2 | 2 | 5 | 35 |
| 171 | SGNSVTQTAAV | activation loop | 1.00 | 1;2 | 2 | 6 | 167 |
| 173 | NSVTQTAAVIG | activation loop | 1.00 | 1;2 | 2 | 5 | 158 |
| 179 | AAVIGTAQYLS | activation loop | 1.00 | 1;2 | 2 | 6 | 27 |
| 294 | TDAERTSLLSS | juxtamembrane segment | 1.00 | 1 | 2 | 2 | 44 |

A. Experimental design for assaying the allosteric activation of PknB by kinase-dead PknB ($PknB^{KD}$).
B. Autoradiograph of a titration experiment. A fixed amount of SNAP-$PknB^{WT}$ was incubated for the same amount of time with increasing amounts of either $PknB^{KD}$ or $PknB^{KD\ L33D}$.
C. Autoradiograph of a titration experiment. A fixed amount of SNAP-$PknB^{WT}$ was incubated for the same amount of time with increasing amounts of either $PknB^{KD}$ or $PknB^{KD\ L33D}$.
D. Autoradiograph of a titration experiment. A fixed amount of SNAP-$PknB^{WT}$ was incubated for the same amount of time with increasing amounts of either $PknB^{KD}$ or $PknB^{KD\ L33D}$.
E. Quantification of the autoradiographs from panels A-C.
F. Intact mass spectrum of SNAP-$PknB^{WT}$ autophosphorylation reaction performed at 20 μM.
G. Tandem mass spectrum following trypsin digest of SNAP-$PknB^{WT}$ autophosphorylation reaction performed at 20 μM.

### Supplementary Methods

#### Molecular dynamics

##### Simulation Parameters

The parent model was based on the inactive IRK crystal structure (PDB code: 1IRK(1)) with ATP and $Mg^{2+}$ introduced into the ATP-binding pocket using a restraining procedure based on the interactions in the active-state of IRK (PDB code: 1IR3)(2). The simulated system contained the 303-residue IRK kinase domain (4,792 protein atoms), ATP (43 atoms), two $Mg^{2+}$ ions, 17,398 TIP4P(3) water molecules, and was charge-neutralized via 9 $K^{+}$ ions, for a total of 74,438 atoms in an 8.3 x 8.3 x 8.3 nm cubic box. After equilibration, for each of the 3 phosphorylatable tyrosine residues in the inactive IRK-ATP-$Mg^{2+}$ system (Y1158, Y1162 and Y1163) a preparatory procedure was applied in order to bring the target tyrosine hydroxyl oxygen to a sub-5 Å distance from the ATP gamma phosphate phosphorus atom (ATP-γP), while keeping the rest of the system structurally unchanged and maintaining ATP and $Mg^{2+}$ in the binding pocket through restraints on the known coordination and binding-pocket interactions(2). The structural integrity and stability of the resulting relaxed, phosphorylation compatible conformations were subsequently assessed using fully unrestrained 1 µs production simulations. All simulations were based on the AMBER99SB-ILDN force field(4), AMBER-compatible ATP parameters(5), a 2 fs time step, Particle Mesh Ewald Electrostatics(6, 7) with 1.2 nm Coulomb and van der Waals cutoffs, and LINCS contraints on bonds to hydrogen(8). The NPT ensemble was equilibrated at T = 300 K with the v-rescale thermostat(9) with a relaxation time $\tau_T$ of 0.1 ps and P = 1 bar with the c-rescale barostat with a relaxation time $\tau_p$ of 2 ps and the compressibility of $k = 4.5e^{-5}$ $bar^{-1}$ (10).

##### Analysis of Phosphorylation-Compatible ATP/OH Distance

The possibility of *cis*-autophosphorylation was assessed by monitoring the distance between ATP-γP and the hydroxyl oxygen of the phosphate acceptor. Using the transition range between associative and dissociative phosphorylation mechanisms(11, 12) as a reference point, together with the detected 5.3 Å distance between substrate tyrosine hydroxyl oxygen and ANP-γP in the active reference structure(2), distances of <=5 Å were considered to be compatible with potential *cis*-autophosphorylation. Interatomic distances were calculated over fully unrestrained trajectories using GROMACS gmx pairdist(13).

#### Root Mean Square Deviation (RMSD) Analysis

RMSD analysis was employed to assess whether ATP-proximal tyrosine conformations required large-scale changes from the inactive kinase structure. Protein trajectories were corrected for periodic-boundary effects, aligned by backbone atoms, and the last 90% compared with the simulation's starting frame as the primary reference using GROMACS gmx rms(13). RMSD was calculated for whole protein backbone and for the non-activation-loop backbone region (981-1146 and 1174-1283). Summary statistics (mean, min, max and standard derivation) were derived from the time-dependent RMSD data.

#### Protein expression, purification, and labeling

Fusion protein constructs were cloned into pFastBac Dual with an N-terminal 10x His tag followed by a GST tag, TEV cleavage site and either the SNAP tag (generation 1) or Halo tag (generation 8) followed by a 3C cleavage site and the insulin receptor kinase (IRK) domain (residues 956-1283). In the case of PknB1, we employed the same construct boundaries of the kinase domain (1-308) previously described(14). The proteins were expressed in baculovirus-infected Sf9 insect cells grown in ESF921 medium (Expression Systems) at 27°C.

Cells were harvested at 4,000 g for 30 min and the pellet was lysed in 100 mL lysis buffer (50 mM HEPES pH 7.4, 150 mM NaCl, 1 mM TCEP, 20 mM imidazole, 2 mM $MgCl_2$, 1x protease inhibitor cocktail (Sigma P8849), 10 nM benzonase (purified in-house)). The lysate was centrifuged at 39,000 g, 4°C for 30 min and the supernatant applied to a HisTrap FF 1 mL (Cytiva). The column was washed with 20 column volumes (CV) of NiA buffer (50 mM HEPES pH 7.4, 150 mM NaCl, 1 mM TCEP) before being eluted in three steps with 5, 10 and 30% NiB buffer (50 mM HEPES pH 7.4, 150 mM NaCl, 1 mM TEP, 1 M imidazole). The 30% NiB elution fractions were pooled, diluted to 100 mM imidazole in NiA buffer and dialyzed against 50 mM HEPES pH 7.4, 100 mM NaCl, 1 mM TCEP overnight at 4 °C. During dialysis, the proteins were dephosphorylated with 10 nmol lambda phosphatase (purified in-house).

The dephosphorylated protein was diluted to 30 mM NaCl with QA buffer (50 mM HEPES pH 7.4, 1 mM TCEP) and loaded onto a 1 mL CaptoQ column (Cytiva) before elution with a linear gradient from 0% to 100% QB buffer (50 mM HEPES pH 7.4, 500 mM NaCl, 1 mM TCEP) in 100 CV. Protein-containing fractions were pooled, concentrated and stored at -80°C. Homogeneous dephosphorylation was verified by intact mass spectrometry.

For protein labeling, single-stranded DNA oligos were purchased from Biomers. For SNAP tag labeling, the oligo was modified with benzylguanine (BG) at its 5' end and fluorescently labeled with Atto488 at its 3' end. For Halo tag labeling, the oligo was modified with PEG2 at its 5' end and fluorescently labeled with Cy5 at its 3' end. To covalently attach the oligo to the protein, a 2.5-fold molar excess of oligo was incubated with protein for 2 h at 4 °C in labeling buffer (50 mM HEPES pH 7.4, 150 mM NaCl, 1 mM TCEP). To remove unbound oligo, the sample was incubated with pre-equilibrated glutathione sepharose beads (Cytiva) in wash buffer (50 mM HEPES pH 7.4, 150 mM NaCl, 1 mM TEP, 0.25% CHAPS) with gentle rotation for 2 h at 4 °C. The beads were washed five times with wash buffer before overnight incubation with 1 nmol TEV (purified in-house) at 4 °C to remove the $His_{10}$-GST tag. Following TEV cleavage, the supernatant was recovered using Mobicol spin columns (MoBiTec) and concentrated before injecting it onto an S75 3.2/300 size exclusion column (Aekta Micro system) equilibrated in storage buffer (50 mM HEPES pH 7.4, 150 mM NaCl, 1 mM TCEP). Protein concentration was estimated by measuring the absorbance of the DNA-protein conjugate at 260 nm. For all steps involving the conjugation oligo (storage, labelling and separation) were performed in DNA LoBind tubes (Eppendorf).

To generate either SNAP-IRK or SNAP-PknB for in-solution kinase assays, the respective GST fusion protein was cleaved with TEV protease overnight at 4 °C. The cleaved sample was incubated with Ni-Sepharose beads (Cytiva) for 30 min at 4 °C in the presence of 10 mM imidazole and the supernatant injected onto a size exclusion chromatography 3.2/300 column (Cytiva) pre-equilibrated in 50 mM HEPES pH 7.4, 150 mM NaCl and 1 mM TCEP.

Kinase-dead IRK ($IRK^{KD}$) was expressed in baculovirus-infected Sf9 cells as a $His_{10}$-GST fusion protein and purified by HisTrap and anion exchange chromatography. The protein was cleaved with TEV protease and dephosphorylated with 10 nmol lambda phosphatase during overnight dialysis into 50 mM HEPES pH 7.4, 100 mM NaCl, 1 mM TCEP, $IRK^{KD}$ was further purified by anion exchange chromatography (CaptoQ) and size exclusion chromatography (Superdex 10/300) in 50 mM HEPES pH 7.4, 150 mM NaCl and 1 mM TCEP.

Both kinase-dead PknB1 constructs (D138N and D138N/L33D) were cloned in a pHis expression vector with an N-terminal 6xHis tag preceded by a TEV cleavage site for expression in BL21 STAR bacterial cells. Cells were grown in TB medium containing 100 μg/mL ampicillin at 37 °C, 200 rpm to an OD = 0.7 before induction with 200 μM IPTG and incubation overnight at 20 °C. The cells were centrifuged at 4,000 g for 30 min and the pellet lysed in 50 mL lysis buffer (50 mM HEPES pH 7.4, 150 mM NaCl, 1 mM TCEP, 10 mM Imidazole, 2 mM $MgCl_2$, 0.2 mg/mL lysozyme, 2 mM benzamidine) per 1 L expression culture. The lysate was sonicated, centrifuged at 39,000 g for 30 min at 4 °C and the supernatant was loaded onto a 5 mL HisTrap column (Cytiva). After washing with 20 mM imidazole (5 CV), the protein was eluted using 300 mM imidazole. The His tag was cleaved off by adding 8.8 nmol TEV protease (purified in house) overnight at 4 °C while simultaneously dialyzing against 50 mM HEPES pH 7.4, 150 mM imidazole and 1 mM TCEP. Cleaved PknB protein was diluted to 50 mM NaCl in Buffer $Q_A$ (50 mM HEPES, pH 7.4, 1 mM TCEP). The protein was loaded onto a CaptoQ anion exchange column (Cytiva). After gradient elution with Buffer $Q_B$ (50 mM HEPES pH 7.4, 500 mM NaCl, 1 mM TCEP) from 0 to 100 % B in 50 CV, the peak fractions were pooled, concentrated and injected onto a S200 10/300 GL size exclusion chromatography column (Cytiva) equilibrated in SEC Buffer (50 mM HEPES pH 7.4, 150 mM NaCl, 1 mM TCEP) and snap frozen.

#### Protein recovery from DNA cylinders

A 12-fold excess of biotinylated oligo complementary to the six overhangs on one side of the cylinders was added for 1 h at room temperature. Streptavidin magnetic beads (ThermoFisher) were equilibrated in bead wash buffer (50 mM HEPES pH 7.4, 150 mM NaCl, 0.25% CHAPS, 10 mM $MgCl_2$) and added to the biotinylated origamis for 1 h at room temperature. The beads were washed extensively with bead wash buffer and the absorbance at 260 nm of the supernatant was monitored on a Nanodrop spectrophotometer until no protein-DNA conjugate was detected. The sample was resuspended at an origami concentration of 100 nM and incubated with the indicated ratios of toe-hold oligo for 2 h at room temperature. The beads were pelleted, the supernatant collected, and the pellet resuspended in an equivalent volume of bead wash buffer before being subjected to densitometry-based analysis by Coomassie-stained SDS-PAGE.

#### Super-resolution microscopy

Super-resolution imaging experiments were performed at room temperature (23C) on a custom-built, epi-illumination widefield and Total Internal Reflection Fluorescence (TIRF) microscope (described in(15)), in TIRF mode. Briefly, the sample was illuminated with a LightHub laser combiner (Omicron-Laserage Laserprodukte) with Luxx 405, 488, 638 and Cobolt 561 nm lasers. In TIRF mode, the light from the laser is coupled into a single mode fiber, and the beam is displaced to the periphery of the objective's back aperture until the TIRF condition is reached, i.e., the background signal originating from fluorescence excitation of the imager strand in solution above the focus plane is minimized. Sample excitation and fluorescence signal was collected through a 160X, 1.43NA oil-immersion objective (Leica), which was focused on the sample using a piezoelectric objective positioner (PIFOC, Physik Instrumente). The fluorescence was recorded with an EMCCD camera (Evolve512D, Photometrics), which also triggers the laser illumination only when recording is ongoing. The sample was kept in the focus position for the duration of the imaging experiments via a closed-loop focus lock feedback system, wherein the light from a 785nm laser (iBeam Smart, Toptica) is reflected by the glass sample support in total internal reflection condition. Thus, any change in focus is translated into a different lateral position of the returning laser light, and the feedback loop acts on the objective positioner to keep the reflection stable, which in turn stabilizes the sample within ± 10nm over hours. Laser triggering and sample stabilization, as well as other essential functions (e.g., data acquisition) were handled via custom electronics (microFPGA), MicroManager 2.0 and the custom-written EMU plugin for micromanager(16–18). To set up an imaging experiment, the sample was initially surveyed using the 488nm laser line at low power and a suitable emission filter, to identify the presence of a sufficient number of DNA origami cylinders in the field of view (FoV). Once a field of view with sufficiently high density of cylinders was identified, a reference image was saved. DNA-PAINT imaging was then performed on the same area, acquiring 50000 frames-long timelapses with an exposure time of 150 ms, illuminating the sample with the 561nm wavelength laser with a power density of approximately 3.5 $kW/cm^2$, and using a suitable emission filter to reject residual excitation light.

#### Super-resolution image reconstruction

The SMLM image reconstruction was performed using SMAP(19), a Matlab-based open source package for superresolution microscopy data analysis available at https://github.com/jries/SMAP. The timelapses originating from the 561nm excitation DNA-PAINT measurement were loaded in the software and the imager strand binding events were detected as maxima in each frame with a difference of Gaussian (DoG) filter. The corresponding fluorophore localizations were estimated by fitting a 2D gaussian model on a 7x7 pixel area around the event, assigning x,y coordinates and calculating variables such as localization precision, number of photons detected and other variables of interest. These localizations are then corrected for lateral drift with an in-built cross correlation-based algorithm in the analysis software. Whenever multiple localization appeared within a user-defined area, they were considered as originating from the same fluorophore and thus grouped together by summing the number of photons detected, and its coordinates recalculated as the photon-count weighted average of the x,y coordinates of all the grouped localizations. Final localizations were then filtered by localization precision (<5nm) and PSFsize (<150nm) to exclude dim localizations. The thus reconstructed super-resolution images were then overlapped with the images of the same FoV obtained with 488nm excitation as a control to verify that the localizations corresponded to a folded DNA origami cylinder.

#### Mass spectrometry

##### Intact protein liquid chromatography-mass spectrometry

LC-MS analysis was performed on a Vanquish™ Horizon UHPLC system (Thermo Scientific) coupled to a Synapt G2-S*i* mass spectrometer (Waters) equipped with a ZSpray ESI source (Waters). Up to 1 pmol of protein in 2 µL solution was loaded onto an XBridge Protein BEH C4 column (300Å, 2.5 µm particle size, dimensions 2.1 mm x 150 mm; Waters) or onto a BioResolve Polyphenyl mAb (450Å, 2.7 µm, 2.1 mm X 150 mm; Waters) with a working temperature of 50°C, 0.1% formic acid (FA) as solvent A, 100% acetonitrile (ACN), 0.08% FA as solvent B. Proteins were separated with a 6 min step gradient from 12% to 72% solvent B (XBridge Protein BEH C4 column) or from 15% to 80% (BioResolve Polyphenyl mAb) at a flow rate of 250 µL/min. The mass spectrometer was operated in resolution mode, with a capillary voltage of 2 kV, a source temperature of 120°C, and a desolvation temperature of 400°C. Data were recorded with MassLynx V 4.2 (Waters) and calibrated by the lockmass of Glu-Fibrinopeptide B. Raw data were analyzed using the MaxEnt 1 process to reconstruct the uncharged average protein mass.

##### Bottom-up liquid chromatography-mass spectrometry

###### *In-gel digestion (ID25207)*

The Coomassie-stained gel bands were destained with a mixture of acetonitrile and 50 mM ammonium bicarbonate (ABC). The proteins were reduced with 10 mM dithiothreitol at 56°C and alkylated with 50 mM iodoacetamide in the dark. Trypsin (Promega; Trypsin Gold, Mass Spectrometry Grade) was used for proteolytic cleavage. Digestion was carried out at 37°C overnight. 10% formic acid (FA) was used to stop digestion and 5% formic acid was used to extract peptides from the gel by sonication.

###### *In-solution digest in sodium deoxycholate buffer (ID26182)*

Protein solution corresponding to ~3 µg was diluted in 50 mM ABC, 2% sodium deoxycholate (SDC). Dithiothreitol (DTT) was added to a final concentration of 10 mM for 30 minutes at 50°C. Reduced thiols were alkylated with 20 mM iodoacetamide for 30 min at RT in the dark. Residual iodoacetamide was quenched with 5 mM DTT. The reaction mixture was diluted to 1% SDC with 50 mM ABC. Proteases (trypsin or chymotrypsin; Promega) were added at a 1:30 protease:protein ratio and incubated at 37°C overnight (trypsin) or at 25°C for 5 h (chymotrypsin). Digestions were stopped by adding 10% TFA to a final concentration of 2%. The precipitated SDC was pelleted by centrifugation at 15000 g for 10 min at RT. Digests were cleaned up using the protocol of Rappsilber et al., 2007 (20) with some modifications. Three C18 Empore disk punches were packed into 200 µL pipette tips. Tips were pre-wetted with

100 µL methanol, followed by 100 µL 80% acetonitrile (ACN), 0,1% trifluoroacetic acid (TFA), and equilibrated with 100 µL 0.1% TFA by centrifugation at 376 g for 5 min. Acidified peptide solutions were applied, and spun at 271 g. Bound peptides were washed with 100 µL 0.1% TFA, and eluted with 2 x 30 µL 40% ACN, 0.1% TFA into PCR tubes. Eluates were dried in a vacuum centrifuge and resuspended in 20 µL 2% ACN, 0.1% TFA.

#### Liquid chromatography-mass spectrometry analysis

LC-MS analysis was performed on a Vanquish Neo UHPLC system (Thermo Scientific) coupled to an Orbitrap Astral mass spectrometer (Thermo Scientific). The system was equipped with a Nanospray Flex ion source (Thermo Scientific), and a Column Heater (IonOpticks) connected to a Heater Controller (IonOpticks).

Peptides were loaded onto a trap column (PepMap Neo C18 5mm × 300 µm, 5 µm particle size, Thermo Scientific) using 0.1% TFA as mobile phase, and separated on an analytical column (Aurora Ultimate XT C18, 25 cm × 75 µm, 1.7 µm particle size, IonOpticks) at 50°C, applying a linear gradient starting with a mobile phase of 98% solvent A (0.1% FA) and 2% solvent B (80% acetonitrile, 0.08% FA), increasing to 35% solvent B over 30 min (60 min) at a flow rate of 300 nL/min.

The Orbitrap Astral mass spectrometer was operated in data-dependent acquisition (DDA) mode with 0.6 s (0.7 s) MS1 cycle time. Survey scans were acquired from 375-2000 m/z, with standard target value for the normalized automatic gain control (AGC), a resolution of 240,000 (120,000) and a maximum injection time of 10 ms. The most intense precursor ions (charge states +2 to +6) were selected for fragmentation using an isolation window of 1.6 m/z. Selected ions were analyzed in the Astral detector with a maximum fill time of 20 ms and standard AGC target after higher-energy collision dissociation (HCD) fragmentation with normalized collision energy of 30%. Monoisotopic precursor selection (MIPS) was set to “peptide” mode, the intensity threshold to 5E3, and selected precursors were dynamically excluded for 10 s (20 s) with isotope exclusion enabled.

#### Data processing

Raw data were analyzed with FragPipe, using MSFragger(21), IonQuant(22) and Philosopher(23). The default FragPipe workflow for label free quantification was used, except ‘Normalize intensity across runs’ and ‘Match between runs’ was turned off. Cleavage specificity was set to the respective protease with 2 missed cleavages allowed. The FDR was set to 1%. Oxidation of methionine, phosphorylation on serine, threonine and tyrosine and N-terminal protein acetylation were specified as variable modifications. Carbamidomethylation of cysteine was set as a fixed modification. MS2 spectra were searched against target sequences in their expression background (Sf9 insect cells) concatenated with a database of common laboratory contaminants (https://github.com/maxperutzlabs-ms/perutz-ms-contaminants). Computational analysis was performed using Python and the Python library MsReport (source code: https://github.com/hollenstein/msreport).
